# Expression of 9-O- and 7,9-O-acetyl modified sialic acid in cells and their effects on influenza viruses

**DOI:** 10.1101/650341

**Authors:** Karen N. Barnard, Brian R. Wasik, Justin R. LaClair, Wendy S. Weichert, Brynn K. Lawrence, Colin R. Parrish

## Abstract

Sialic acids (Sia) are widely displayed on the surfaces of cells and tissues. Sia come in a variety of chemically modified forms, including those with acetyl modifications at the C7, C8, and C9 positions. Here, we analyzed the distribution and amounts of these acetyl modifications in different human and canine cells. As Sia or their variant forms are receptors for influenza A and influenza C viruses, we examined the effects of these modifications on virus infections. We confirmed that 9-*O*-acetyl and 7,9-*O*-acetyl modified Sia are widely but variably expressed across cell lines from both humans and canines. While they were expressed on the cell surface of canine MDCK cell lines, they were located primarily within the Golgi compartment of human HEK-293 and A549 cells. The *O*-acetyl modified Sia were expressed at low levels of 1-2% of total Sia in these cell lines. We knocked out and over-expressed the sialate *O*-acetyltransferase gene (CasD1), and knocked out the sialate *O*-acetylesterase gene (SIAE) using CRISPR/Cas9 editing. Knocking out CasD1 removed 7,9-*O*- and 9-*O*-acetyl Sia expression, confirming previous reports. However, over-expression of CasD1 and knockout of SIAE gave only modest increases in 9-*O*-acetyl levels in cells and no change in 7,9-*O*-acetyl levels, indicating that there are complex regulations of these modifications. These modifications were essential for influenza C infection, but had no obvious effect on influenza A infection.

**IMPORTANCE:** Sialic acids are key glycans that are involved in many different normal cellular functions, as well as being receptors for many pathogens. However, Sia come in diverse chemically modified forms. Here we examined and manipulated the expression of 7,9-*O*- and 9-*O*-acetyl modified Sia on cells commonly used in influenza virus and other research by engineering the enzymes that produce or remove the acetyl groups.

## INTRODUCTION

Sialic acids (Sia) are a family of nine-carbon monosaccharides expressed mainly in vertebrates that serve as terminal residues of carbohydrate chains on cell membrane glycoproteins and glycolipids, as well as on secreted glycoproteins at all mucosal surfaces (**Fig 1A**) (1, 2). Sia are key mediators of many cell and tissue functions, where they are bound by cellular receptors such as selectins and siglecs (3, 4). Their ubiquitous presence on cells, tissues, and mucosal surfaces also make Sia a key point of contact for commensal microbes and invading pathogens, including viruses, bacteria, and parasites (3).

**Figure 1.**
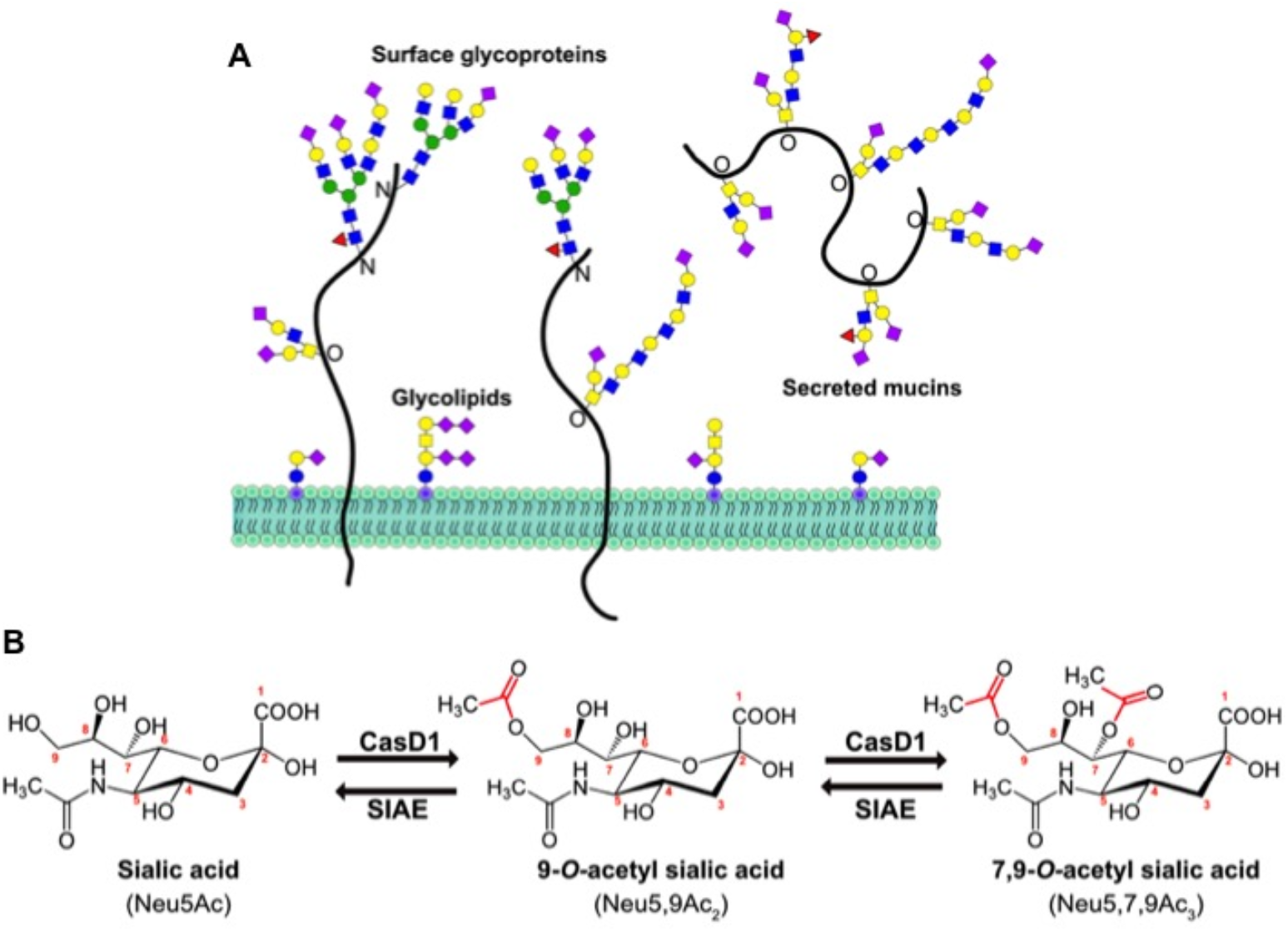
(A) Sialic acids (purple diamonds) terminate glycan chains on glycolipids and glycoproteins as part of the glycocalyx on the surface of cells. They can also terminate glycans on secreted glycoproteins, like mucins, that make up the protective mucosal barrier in gastrointestinal and respiratory tissue. (B) The sialate *O*-acetyltransferase, CasD1, adds acetyl groups to sialic acid (N-acetylneuraminic acid, Neu5Ac) at the C-7 from which it migrates to the C-9 position (Neu5,9Ac_2_) under physiological conditions. This can allow for an additional acetyl group to be added by CasD1 to C-7 (Neu5,7,9Ac_3_). The sialate *O*-acetylesterase, SIAE, can remove these acetyl modifications, restoring the unmodified Neu5Ac form of sialic acid.

Sia are highly diverse as there are more than 50 different chemically distinct variants that are formed from the basic structure of N-acetylneuraminic acid (Neu5Ac) by the addition of chemical groups at various positions on the pyranose ring or the glycerol side chain. These modifications include acetyl, sulfo, methyl, and lactyl groups, among others (1). As many different enzymes and pathways introduce these modifications, there are often complex mixtures of Sia forms, with significant variation in both the levels and specific combinations of each modification (1, 2, 5).

Common chemical additions include *O*-acetyl modifications to the C-4, 7, 8, and/or 9 positions, resulting in a variety of combinations including Neu4,5Ac, Neu5,9Ac_2_, Neu5,7,9Ac_3_ Sia. The addition of *O*-acetyl modifications to the C7 and/or C9 positions is mediated by the sialate *O*-acetyltransferase enzyme, Cas1 domain containing 1 (CasD1) (**Fig. 1B**). CasD1 appears to add acetyl groups to the C7 position, from which it has been observed to migrate to the C8 and C9 position under physiological conditions, allowing the possible addition of another acetyl group to C7 (6, 7). A migrase enzyme has been proposed to aid in the transfer of the acetyl group from C7 to C9, however a specific enzyme has yet to be identified (5, 8, 9). CasD1 is localized in the late Golgi, so acetyl modifications are added during the later stages of protein glycosylation. The regulatory processes that control the number of acetyl groups added or their positions have not been well defined, although clear differences in expression of 7,9-*O*-Ac and 9-*O*-Ac have been reported in mouse and human tissues, chicken embryos, and some other animals (10, 11). CasD1 uses acetyl-CoA and likely has a preference for CMP-Neu5Ac as a substrate, so that it is less active on CMP-Neu5Gc (7). The existence of an enzyme to facilitate the migration of acetyl groups from C7 has been proposed, but not identified (5).

At least one sialate O-acetylesterase (SIAE) enzyme regulates the display of 7,9-*O*- and 9-*O*-Ac in many cells and tissues (**Fig. 1B**). The SIAE gene encodes two isoforms that vary in the presence of a proposed C-terminal localization tag in the lysosomal (Lse) form that is absent from the cytosolic form (Cse) (12). However, the processes that regulate the expression and activity of these two isoforms remain poorly defined. Lse has been found to localize to the Golgi and/or ER and on the surface of cells when over-expressed, with the majority being secreted into the supernatant (12). Antibody staining shows that Cse is found diffusely throughout the cytosol, where it is thought to remove 9-*O*- and 7-*O*-acetyl groups to recycle the Sia for reuse in glycosylation (12). Despite the reports of distinct protein expression in mouse tissues, bioinformatic analysis of RNA expression in human cells and tissues show mRNA corresponding to the Lse form of SIAE and none responding to the Cse form (13). It is therefore still unclear how these two isoforms are regulated in humans or other animals, whether their expression relates to CasD1, or what controls the levels and locations of 9-*O*- and 7,9-*O*-Ac expression.

Sia *O*-acetylation and de-acetylation play important roles in many different biological processes, particularly in development, cancer, and immunology. For example, *O*-acetyl modifications to Sia may alter the binding of host lectins, including the Sialic acid-binding immunoglobulin-type lectins (Siglecs) (3, 5, 14). Siglecs are regulators of many different cell functions and developmental processes – examples include B- and T-cells where the presence of 9-*O*-acetyl Sia modulates immune cell activation and differentiation (15). In B-cells, negative regulation of B-cell receptor (BCR) activation by Siglec CD22 is mediated by binding to Neu5Ac-terminated glycan chains on the BCR, which can be blocked by *O*-acetyl modifications (16, 17). The presence of 9-*O*-Ac can also reduce the activity of sialidases, including human neuraminidases (18). Incorrect regulation of 9-*O*-Ac, 7,9-*O*-Ac, and SIAE activity has been linked to autoimmune disorders through the development of auto-antibodies (17, 19). 9-*O*-Ac and 7,9-*O*-Ac and their regulation by SIAE also appear to play important roles during early stages of embryonic development, spermatogenesis, and in different forms of cancer including acute lymphoblastic leukemia, colon cancer, and breast cancer (20–23).

Effects of different Sia modifications have also been suggested for the binding of pathogens or on the activities of their sialidases (neuraminidases). However, in general these are still not well documented, with the exception of those that use the modified forms as receptors. Both influenza A (IAV) and influenza C (ICV) viruses use Sia as their primary receptors for host recognition and cell entry, but with different effects of Sia modification. IAV interacts with Sia through two surface glycoproteins, hemagglutinin (HA) and neuraminidase (NA). HA binds Sia to initiate the endocytic uptake of the virus by the cell, leading to fusion between the viral envelope and the endosomal membrane at low pH (24). NA is a sialidase which cleaves Sia off glycan chains when it is present in mucus or on the surface of cells, allowing the virus to penetrate to the epithelial cells, and also preventing aggregation of newly produced virus after budding (25, 26). In contrast to IAV, ICV has one surface glycoprotein, the hemagglutinin-esterase fusion protein (HEF), which serves similar purposes to both HA and NA. HEF binds specifically to 9-*O*-acetyl Sia to initiate uptake of the virus into cells, while the esterase domain removes 9-*O*-acetyl modifications, releasing the virus from mucus, and disassembling virus aggregates after budding (27). While the role of *O*-acetyl modified Sia for ICV infection is well documented, how these modifications affect IAV is not well characterized. Previous limited studies have suggested that the presence of 9-*O*-Ac on cells may be inhibitory for NA activity and HA binding of IAV (28, 29).

Here we examine and more closely define the expression and distribution of 9-O-acetyl and 7,9-O-acetyl Sia on cells in culture, define the effects of SIAE and CasD1 on display of these Sia modifications, and perform an initial examination of the effects of these modified Sia on infection by IAV and ICV. We used CRISPR-Cas9 for gene engineering of the enzymes that add or remove the 9-*O*- and 7,9-*O*-acetyl groups from Sia, and combine these with the recently developed viral protein-derived probes that specifically recognize modified Sia. By merging these with HPLC-based methods for quantifying the different Sia forms, we provide a more detailed understanding of the expression and localization of these modifications in cells, and examine their affects on two host-virus interactions as examples.

## RESULTS

### Expression of 7,9-*O*- and 9-*O*-acetyl Sia in cells

There is currently only sporadic information about the expression of modified Sia on commonly used cell lines, or an understanding of how that compares to the expression in animal tissues. We examined cell lines that are widely used in many experimental systems: A549 human type II alveolar epithelial cells, HEK-293 human kidney derived cells (possibly embryonic adrenal precursor cells (30)), and MDCK canine kidney epithelial cells. Additionally, we tested MDCK-type I and type II cells that were previously sub-cloned from the ATCC MDCK line (MDCK-NBL2) and have been extensively characterized (31–33). We used probes derived from porcine torovirus (PToV) and bovine coronavirus (BCoV) hemagglutinin-esterase proteins (HE) that were fused to human IgG1 Fc and had the esterase active site inactivated (HE-Fc). The PToV HE-Fc probe recognizes 9-*O*-Ac, while BCoV HE-Fc recognizes primarily 7,9-*O*-Ac but also has a low affinity for 9-*O*-Ac (10, 11). By immunofluorescence microscopy, the different forms were present at variable levels, with 10 to 70% of the cells showing staining under standard culture conditions (**Fig. 2A,B**). MDCK-NBL2 and MDCK-type I cells showed both strong surface and internal staining for 7,9-*O*- and 9-*O*-Ac forms, while both were mostly found in intracellular locations in A549 and HEK-293 cells, with an occasional cell showing bright surface staining. MDCK-type II cells showed staining only for 9-*O*-Ac and none for 7,9-*O*-Ac, indicating that these modifications are regulated independently. In HEK-293 and A549 cells, both 7,9-*O*- and 9-*O*-Ac appeared to be localized within the Golgi compartment as confirmed by co-staining with the Golgi marker GM130 (**Fig. 3**). Similar localization differences between cell lines have been seen previously using the ICV HEF as a probe (34). In many of the cell lines, there was inherent variability within populations in terms of level of staining and localization. For example, in MDCK-NBL2 not all cells were positive for 9-*O*-Ac, while in MDCK-type I some cells retained more 9-*O*-Ac and 7,9-*O*-Ac internally and others displayed more of these modified Sia on their surface (**Fig. 2A**). This heterogeneity was consistent between different passages of each cell line.

**Figure 2.**
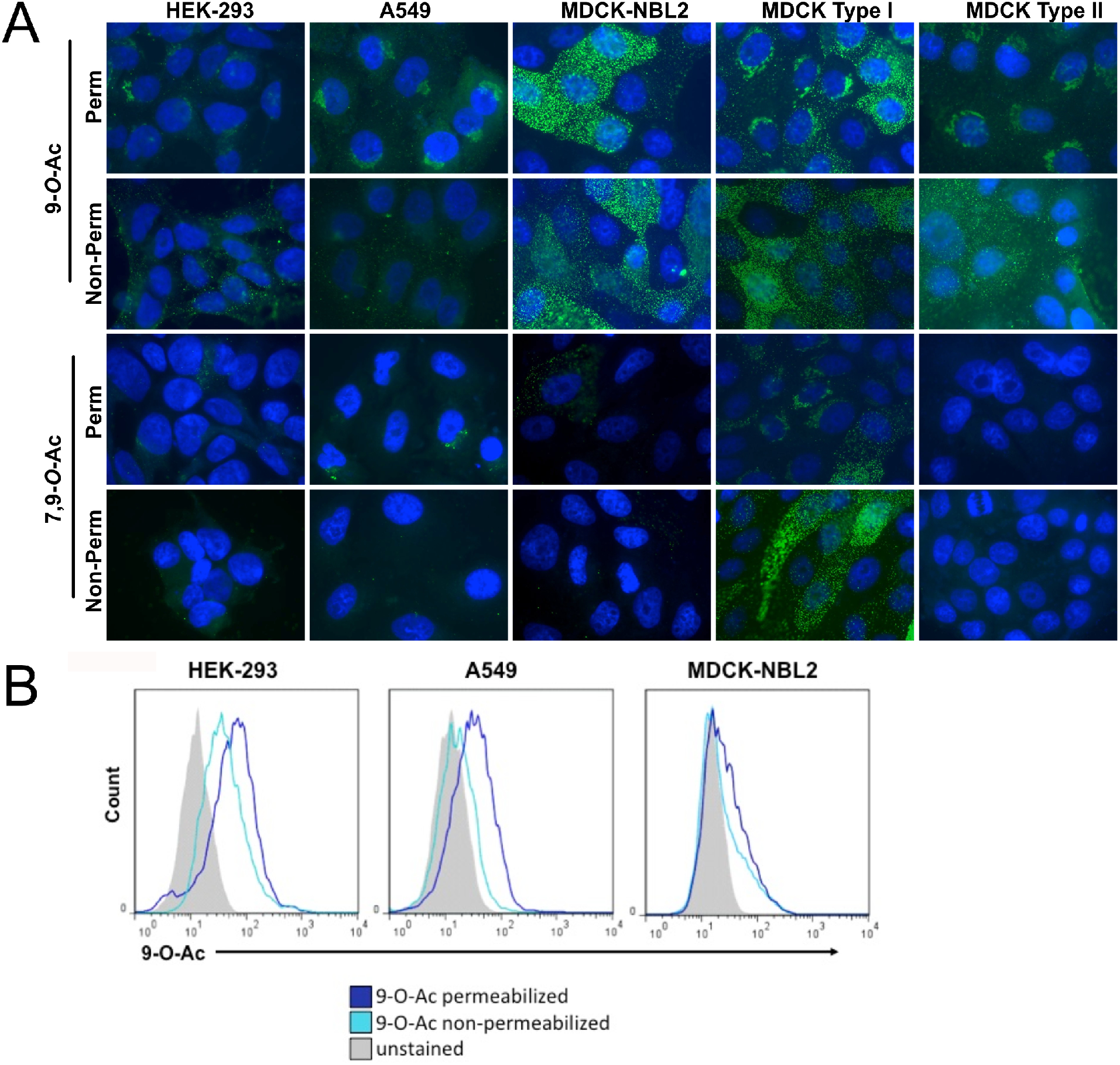
Surface and internal expression of 9-*O*-Ac and 7,9-*O*-Ac on different cell lines. (A) Fluorescent staining of human HEK-293, A549, and canine MDCK AATC line (NBL2), MDCK type I, and MDCK type II. Cells were probed with HE-Fcs probes derived from BCoV and PToV, which recognize 9-*O*-Ac and 7,9-*O*-Ac respectively. Cells were permeabilized (perm) using Carbo-Free blocking reagent with 0.001% Tween-20 while non-permeabilized cells (non-perm) received only Carbo-Free block. All cells imaged at 60 ×, nuclei stained with DAPI. (B) Representative flow cytometry graphs showing distribution of positive staining for HEK-293, A549 and MDCK-NBL2 cell lines. BCoV and PToV HE-Fcs probes were used and permeabilization (perm) and non-permeabilization (non-perm) methods were as in IFA staining.

**Figure 3.**
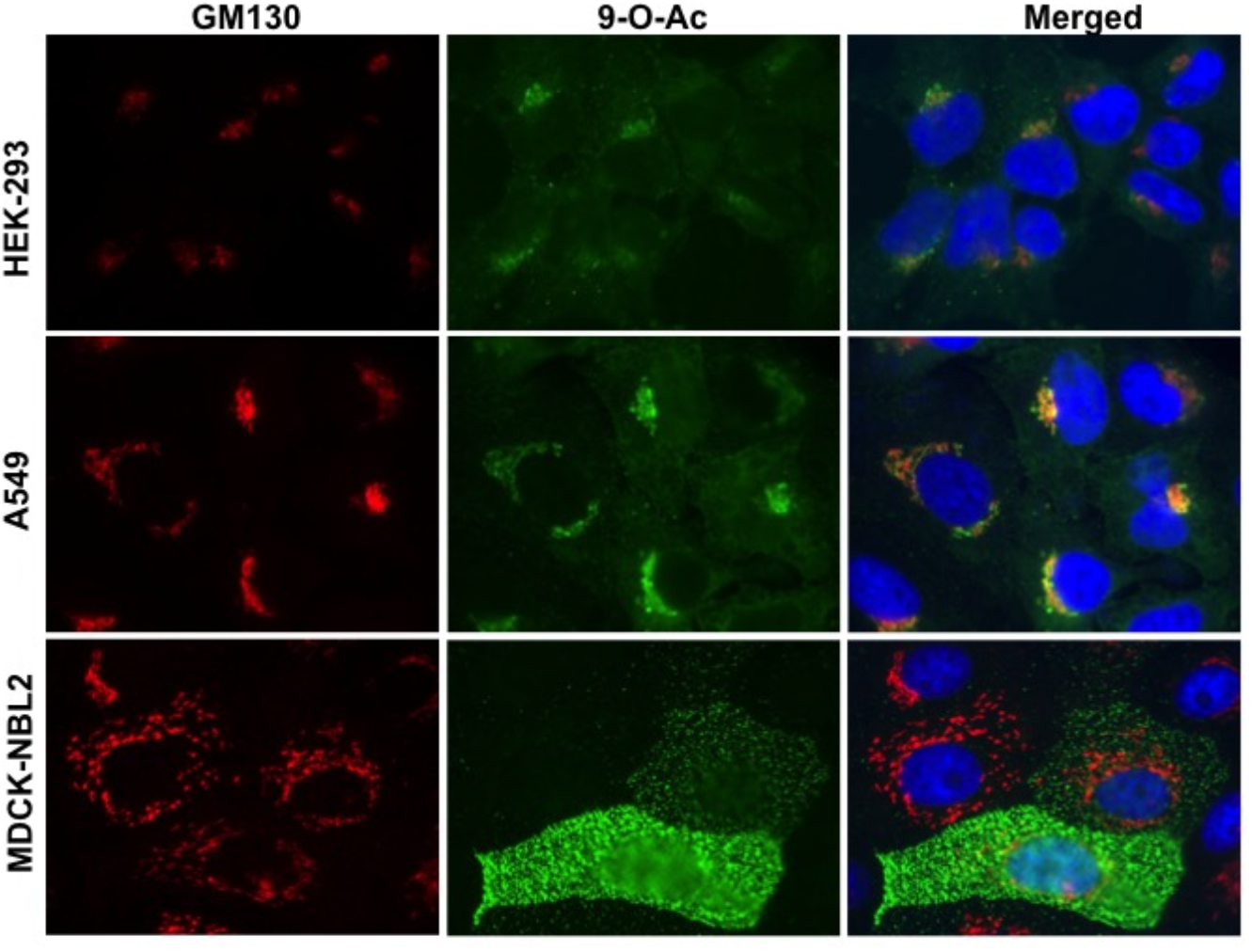
Staining of HEK-293, A549, and MDCK-NBL2 cells with PToV HE-Fc for 9-O-Ac, and co-staining for the Golgi marker GM130. Cells were permeabilized using 0.001%Tween-20 and imaged at 60x magnification.

The expression levels of those modified Sia variants were also quantified by HPLC analysis using DMB labeling and fluorescence detection (35). The cells were treated at 80°C for 3h in the presence of 2M acetic acid, and likely represent the total Sia present in the cells. Even though some cell lines showed strong staining by the HE-Fc probes, levels of 9-*O*-Ac were 1-2% of the total collected Sia from all cells while 7,9-*O*-Ac was not detected in any cell lines (**Fig. 2; Table 1; Supplemental Fig. 3**). This includes cells from other species including cat, mouse, horse, and swine. Unique among the cells tested, A549 cells are secretory and express significant levels of mucin (including MUC1 and MUC5B) (36). We found that A549 cells were able to secrete MUC5B into conditioned media (**Supplemental Fig. 1A**), although we found that the amount secreted was inconsistent between collections. Analysis of conditioned media from A549 cells showed approximately 2% of Sia was 9-*O*-Ac on secreted proteins with no 7,9-*O*-Ac detected, indicating that at least in these cells, secreted proteins were not enriched for *O*-acetylated Sia (**Supplemental Fig. 1B**). However, it would be worth examining mucins secreted by primary cells or that are present on tissue, to confirm that the secreted proteins from A549 are representative of respiratory mucus.

The low levels of 9-*O*- and 7,9-*O*-Ac detected on HEK-293, A549, and MDCK cells by HPLC could be due to the heterogeneity of the population, as this method measures total Sia for all cells in the population. However, it is likely that even on higher expressing cells, 7,9-*O*- and 9-*O*-Ac make up a small proportion of the total Sia present in the cell or glycocalyx. Previous research has indicated that podoplanin (GP40 in canine cells) is a primary carrier of 9-*O*-Ac on MDCK cells and therefore acts as the main receptor for ICV (37, 38). However, when we co-stained MDCK cells with an anti-podoplanin antibody and PToV HE-Fc, we saw little correlation between podoplanin and 9-*O*-Ac staining (**Supplemental Fig. 2**).

**Table 1.** Total sialic acids were collected from different cell lines via mild acid hydrolysis and analyzed using HPLC. Percentages are out of total sialic acid collected. Representative chromatograms for standard, HEK-293, A549, and MDCK-NBL2 are presented in **Supplementary Figure 3**. n/d = not detected. * primary swine nasal (SiNEC) and tracheal (SiTEC) epithelial cells, courtesy of Dr. Stacey Schultz-Cherry.

| Cell Type | Species | %Neu5Gc | % Neu5Ac | % 9-O-Ac | % 7,9-O-Ac |
| --- | --- | --- | --- | --- | --- |
| A549 | Human | 2.21 | 96.40 | 1.39 | n/d |
| HEK-293 | Human | 2.26 | 96.72 | 1.02 | n/d |
| MDCK-NBL2 | Canine | 0.79 | 97.89 | 1.32 | n/d |
| MDCK Type I | Canine | 0.66 | 97.67 | 1.67 | n/d |
| MDCK Type II | Canine | 0.76 | 98.13 | 1.11 | n/d |
| NLFK | Cat | 2.00 | 96.87 | 1.13 | n/d |
| EqKc3 | Horse | 2.93 | 97.07 | n/d | n/d |
| NBL6 | Horse | 2.52 | 96.38 | 1.1 | n/d |
| L cells | Mouse | 3.87 | 96.13 | n/d | n/d |
| A72 | Canine | 2.04 | 95.85 | 2.11 | n/d |
| SiNEC | Swine* | 4.69 | 93.56 | 1.75 | n/d |
| SiTEC | Swine* | 1.97 | 96.54 | 1.49 | n/d |

\*primary nasal and trachea epithelial cells, courtesy of Dr. Schultz-Cherry

### Production of CasD1 knock out and over-expressing cells

#### Knock out and over-expression of CasD1

To better understand the control of expression of 7,9-*O*- and 9-*O*-Ac, glyco-engineered cells lines were created by manipulating the expression of CasD1 and SIAE genes. To do this, we knocked out CasD1 via CRISPR-Cas9 editing, or over-expressed CasD1 via transfection of an expression plasmid. Knock-out variants of CasD1 (ΔCasD1) were prepared from MDCK-NBL2, A549, and HEK-293 cells using CRISPR-Cas9 targeting of early exons in the CasD1 gene (**Fig. 4A**). ΔCasD1 clones were confirmed by examining for 7,9-*O*- and 9-*O*-Ac display, and then by PCR and sequencing of the genomic region surrounding the deletion in modified Sia negative clones (**Fig. 4A**). For all cells types, ΔCasD1 variants showed loss of both 7,9-*O*- and 9-*O*-Ac display when probed with the different specific HE-Fc probes via flow cytometry and immunofluorescence microscopy (**Fig. 4B, C, D**). This agrees with previous findings that loss of CasD1 leads to loss of both 7,9-*O*- and 9-*O*-Ac modifications in haploid HAP1 cells (7). HPLC analysis of total Sia showed a significant decrease in 9-*O*-Ac expression compared to wild-type (WT) cells, however very low levels (<1%) were still detectable, despite a lack of staining with the HE-Fc probes (**Fig. 4E**). This could be due to exogenous Sia from the fetal bovine serum in the growth media, as is the case for Neu5Gc which is also detectable at similarly low levels on cells even though the gene for Neu5Gc synthesis, CMAH, is not functional in humans or canines (**Supplemental Fig. 3**) (39, 40).

**Figure 4.**
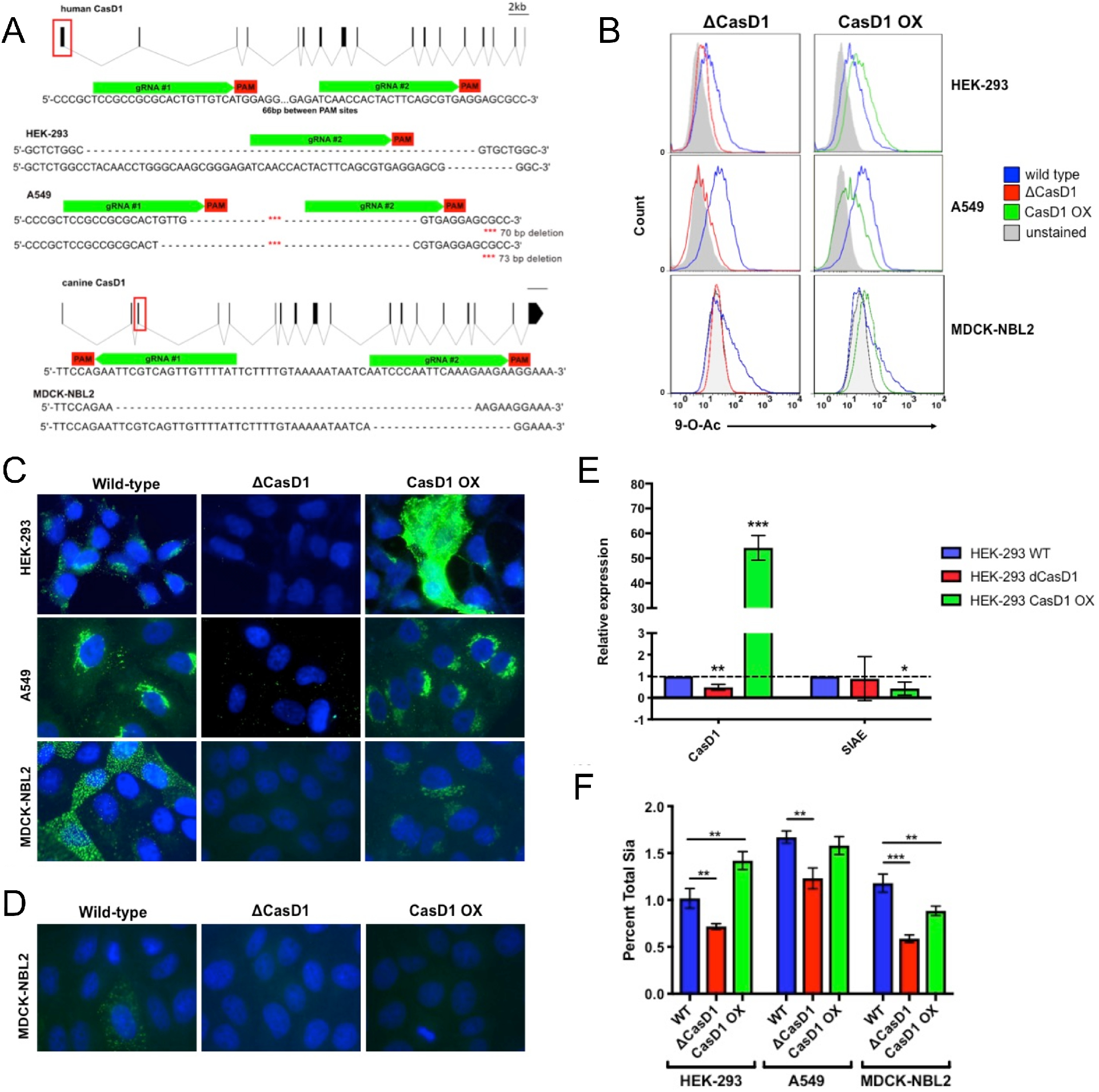
Editing expression of CasD1 in A549, HEK-293, MDCK-NLB2 cells. (A) Schematic of edits in the CasD1 gene and genotypes of edited cells for HEK-293, A549, and MDCK-NBL2 cells. (B) Phenotype of edited cells by flow cytometry using 9-*O*-Ac probe (PToV HE-Fc). Cells were permeabilized using 0.001% Tween-20 to determine internal expression, as most modified Sia is retained internally. The graph is representative of three independent experiments. (C) Immunofluorescence microscopy images of the different engineered cells stained with PToV HE-Fc to detect 9-*O*-Ac. Cells were permeabilized to reveal both surface and internal expression. Cells imaged at 60× magnification. (D) Staining of MDCK WT, ΔCasD1, and CasD1-OX showing representative display of 7,9-*O*-Ac using the BCoV HE-Fc probe. Cells were permeabilized to reveal both surface and internal expression. Cells imaged at 60× magnification. (E) qPCR of CasD1 and SIAE expression in HEK-293 WT, ΔCasD1, CasD1-OX cells. CasD1 still shows mRNA due to mismatch between qPCR primers and edit sites, mRNA is still present but doesn’t produce functional protein. Expression relative to house-keeping gene, GAPDH. Data analyzed using t-test in PRISM software. * = p-value ≤0.05; ** = p-value ≤0.01; *** = p-value ≤0.001. (F) HPLC data for total Sia collected from cells by mild acid hydrolysis for WT, ΔCasD1, and CasD1 OX in HEK-293, A549, and MDCK cells. Data analyzed by t-test using PRISM software. * = p-value ≤0.05; ** = p-value ≤0.01; *** = p-value ≤0.001.

Due to the heterogeneity and low expression of 7,9-*O*- and 9-*O*-Ac in WT cells, we sought to engineer cells with more homogeneous and higher levels of these modifications. We over-expressed the human CasD1 (CasD1-OX) in the ΔCasD1 variants of MDCK-NBL2, A549, and HEK-293 cells, expecting that expression of CasD1 under a strong promoter in the ΔCasD1 background would increase the consistency of the synthesis and display of 9-*O*- and 7,9-*O*-acetyl Sia relative to WT. CasD1-OX cells showed significantly higher levels of CasD1 mRNA compared to WT cells, indicating that the CasD1 expression plasmid was being transcribed at high levels (**Fig. 4E**). However, only a modest increase in average 9-*O*- and 7,9-*O*-Ac expression across the population in HEK-293 cells was seen by flow cytometry, while fluorescent microscopy showed heterogeneity of expression in the population (**Fig. 4B, C, D**). The transfected MDCK and A549 CasD1-OX cells showed recovery of 9-*O*- and 7,9-*O*-Ac synthesis when analyzed by flow cytometry and immunofluorescence microscopy, but levels were not as high as seen in WT cells (**Fig. 4B, C, D**). HPLC analysis confirmed the expression levels for HEK-293, A549, and MDCK cells (**Fig. 4F**). Additionally, heterogeneity was still seen in these populations for 9-*O*-Ac. This suggests that 7,9-*O*- and 9-*O*-Ac expression is not directly regulated by the levels of CasD1 gene expression, but may be affected by post-translational regulation of CasD1 activity or removal of the modifications by SIAE or other enzymes. In addition, these modifications are not regulated the same across individual cells as evidenced by the population heterogeneity. SIAE transcripts were seen to be consistently expressed in HEK-293 WT, ΔCasD1, and CasD1-OX cell populations by qPCR analysis, although SIAE mRNA levels in CasD1-OX cells were lower than those seen in WT (**Fig. 4E**).

#### SIAE knock out cells and display of modified Sia

To determine the role of SIAE activity in regulating 7,9-*O*- and 9-*O*-Ac display, SIAE knockout (ΔSIAE) cells were generated from WT HEK-293 and A549 cells by targeting early exons in the SIAE gene (**Fig. 5A**). Complete knockout of SIAE was confirmed both by genotyping to show deletions in both alleles and by loss of mRNA by qPCR (**Fig. 5A, D**). ΔSIAE cells showed a small but significant increase in surface display and a large increase in internal display of 9-*O*-Ac based on flow cytometry, but did not appear to show any changes in surface or internal display of 7,9-*O*-Ac (**Fig. 5B, C**). HPLC analysis also showed a small increase of 9-*O*-Ac in HEK-293 cells, but no 7,9-*O*-Ac was detected in either cell line (**Fig. 5E**). When the overexpression CasD1 plasmid was transfected into ΔSIAE cells (ΔSIAE+CasD1), cells showed an increase in surface and internal 9-*O*-Ac, but no increase in surface or internal display of 7,9-*O*-Ac by either flow cytometry or immunofluorescence microscopy (**Fig. 5B,C**). HPLC confirmed the flow cytometry results by showing a small increase of 9-*O*-Ac levels in HEK-293 cells that were similar to those seen in CasD1-OX (**Figs. 5E**). A549 cells showed a small but non-significant increase in 9-*O*-Ac levels compared to WT by HPLC analysis. Interestingly, HEK-293 ΔSIAE+CasD1 cells grew more slowly than either ΔSIAE or WT cells (**Fig. 5F**). However, A549 ΔSIAE+CasD1 showed increased growth rates compared to ΔSIAE or WT cells, while ΔSIAE cells had slightly lower growth rates compared to WT. This suggests that 7,9-*O*- and 9-*O*-Ac may affect cell metabolism and growth rates, and that these effects may be cell type specific. Overall, these results show that SIAE regulates levels of 9-*O*-Ac and 7,9-*O*-Ac, but that knocking out SIAE only leads to small increases in these modifications. However, there appear to be mechanisms regulating the surface display of 9-*O*-Ac and 7,9-*O*-Ac, as most of the modified Sia were specifically retained in the Golgi on human HEK-293 and A549 cells, in comparison to WT MDCK cells which appear to display most modified Sia on their surface.

**Figure 5.**
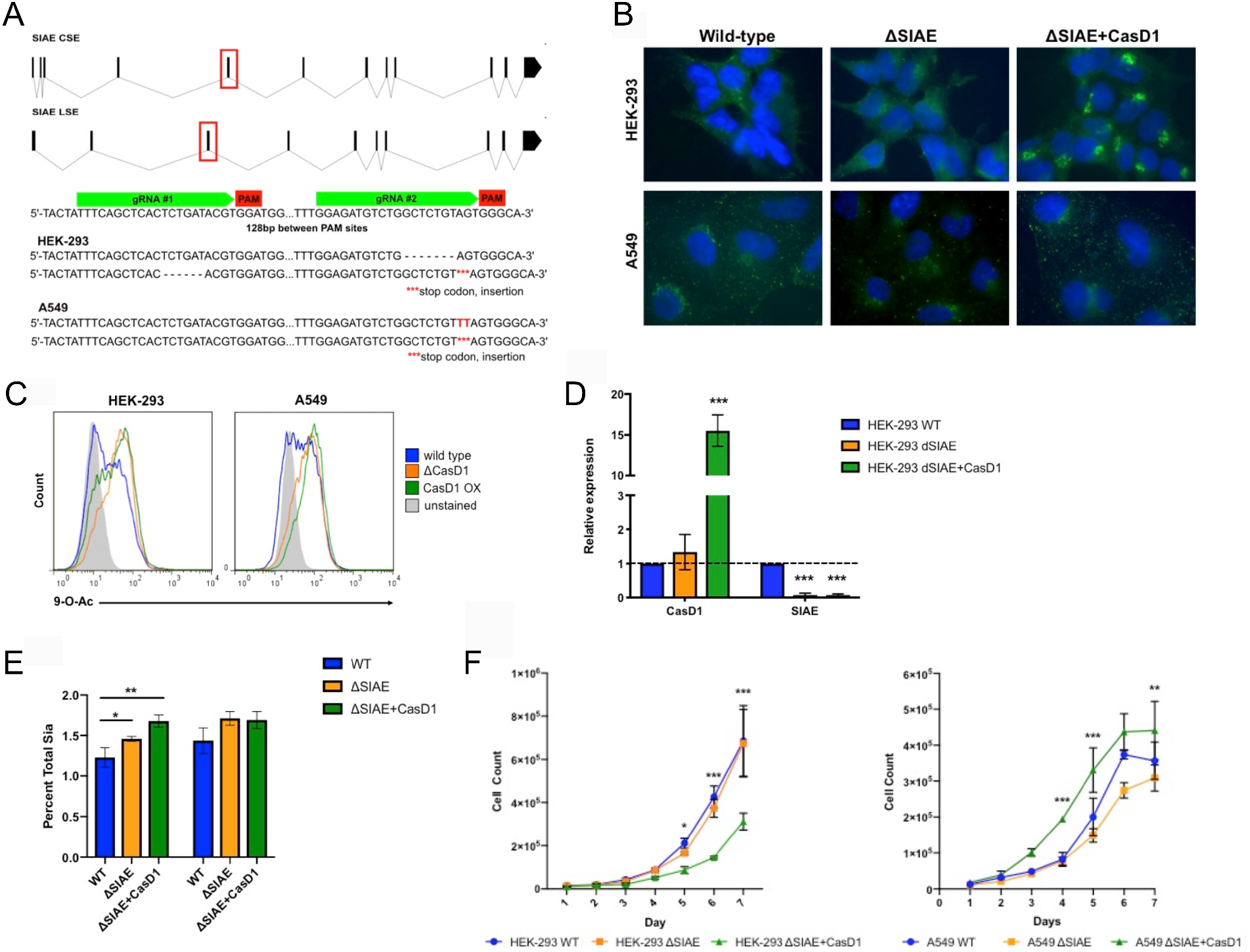
Editing the expression of SIAE in HEK-293 and A549 cells. (A) Schematic of edits in the SIAE gene, which was targeted to remove both isotypes of SIAE. The genotypes of edited cells show frame-shifts that lead to stop codons in all cases. (B) Staining of the different engineered cells with PToV HE-Fc to detect 9-*O*-Ac. Cells were permeabilized using 0.001% tween-20 to determine surface and internal expression. (C) Flow cytometry using PToV HE-Fc showing relative display of 9-*O*-Ac. Cells were permeabilized using 0.001% tween-20 to show both surface and internal expression. Graph is representative of three independent experiments. (D) qPCR of relative SIAE and CasD1 mRNA expression in HEK-293 WT, ΔSIAE, and ΔSIAE+CasD1 cells compared to GAPDH. SIAE qPCR primers overlap with edit site. Data analyzed by t-test using PRISM software. * = p-value ≤0.05; ** = p-value ≤0.01; *** = p-value ≤0.001. (E) HPLC data for total Sia collected from cells by mild acid hydrolysis for WT, ΔSIAE, and ΔSIAE+CasD1 in HEK-293 and A549 cells. Data analyzed by t-test using PRISM software. *= p-value ≤0.05; ** = p-value ≤0.01; *** = p-value ≤0.001. (F) Growth curve WT, ΔSIAE, and ΔSIAE+CasD1 in HEK-293 and A549 cells. Cells were counted every 24 hours. Experiment was performed in triplicate. Data analyzed by 2-way Anova using PRISM software.* = p-value ≤0.05; ** = p-value ≤0.01; *** = p-value ≤0.001.

#### Effects of 7,9-*O*-Ac and 9-*O*-Ac on influenza A and influenza C virus infection

To test the effects of 9-*O*-Ac and 7,9-*O*-Ac on virus infection, WT, ΔCasD1, and CasD1-OX HEK-293 cells were inoculated with human H1N1 (A/California/04/2009) and human H3N2 (A/Victoria/361/2011) IAV strains, and found no significant difference in infection efficiency in any of these cells (**Fig. 6A**). IAV strains also showed equal infection efficiency on MDCK WT, ΔCasD1, and CasD1-OX (**Fig. 6C**). However, ICV strains C/Ann Arbor/1/50 and C/Taylor/1233/1947 showed no infectivity in MDCK ΔCasD1 (**Fig. 6B**). Only the Ann Arbor strain recovered infectivity in MDCK CasD1-OX at much lower levels than in MDCK WT, likely due to the lower surface expression of 9-*O*- and 7,9-*O*-Ac in these cells. It therefore appears that 7,9-*O*-Ac and 9-*O*-Ac do not strongly influence IAV infection, likely due to their low levels of expression. However, viruses such as ICV are able to utilize these low levels of modified Sia as their primary receptors for binding and infection, but cannot infect when they are removed.

**Figure 6.**
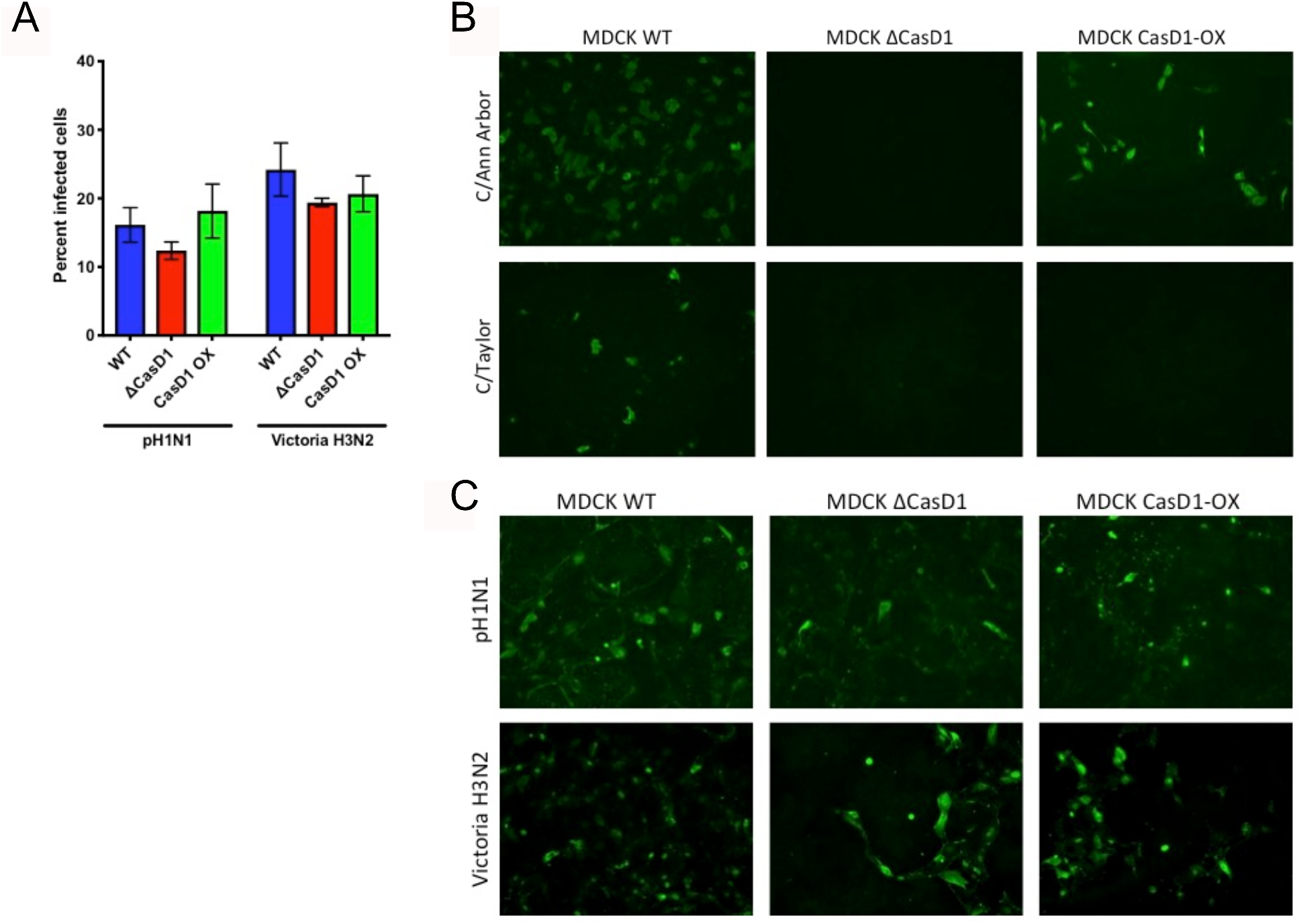
Infection of WT, ΔCasD1, and CasD1-OX cells by IAV and ICV. (A) HEK-293 WT, ΔCasD1, CasD1-OX cells were inoculated at MOI 0.1 with IAV strains pH1N1 (A/California/04/2009) and Victoria H3N2 (A/Victoria/361/2011). Cells were fixed at 24 hours and infected cells per field were counted. Experiment was performed in triplicate. Data analyzed by 2-way Anova using PRISM software. (B) MDCK WT, ΔCasD1, and CasD1-OX cells were inoculated at high MOI with ICV strains C/Ann Arbor/1/50 and C/Taylor/1233/1947 for 48 hr, then imaged. Representative image of three independent experiments. (C) MDCK WT, ΔCasD1, and CasD1-OX cells were inoculated at high MOI with IAV strains pH1N1 (A/California/04/2009) and Victoria H3N2 (A/Victoria/361/2011) for 48 hr, then imaged. Representative image of three independent experiments.

## DISCUSSION

9-*O*-Ac and 7,9-*O*-Ac Sia are widely expressed within tissues and on mucosal surfaces of many animals, but with significant variation in the amounts in different cells, tissues, and animals (10, 11). These modified Sia are present in secreted mucus on mucosal surfaces, including GI and respiratory tissues, where they can potentially control many interactions with both normal flora and pathogens. However, there is still little known about the details of their display levels, cell association, and the ways in which their synthesis is regulated either in cells or on mucosal surfaces. While there have been suggestions that 9-*O*- and 7,9-*O*-Ac might influence IAV infection by interfering with HA binding or NA activities, direct evidence for their effects is sparse. In contrast, ICV is known to use 9-*O*-Ac as its primary receptor for cell binding and infection. Here we use a number of new tools to define the cell-specific expression of 9-*O*- and 7,9-*O*-Ac and provide a preliminary test of their effects on IAV and ICV infection.

We confirmed that 9-*O*- and 7,9-*O*-Ac are expressed on cells in culture and that expression varies between cell lines, as has been previously reported (34). However, when present these modified Sia made up only 1 to 2% of the total Sia when analyzed by HPLC. Cells showed distinct population heterogeneity in 7,9-O- and 9-*O*-Ac display and localization that was observed over many passages examined. Previous studies using ICV HEF probes found similar staining patterns on some cell lines and also showed that after sorting into high and low staining populations, both populations returned to previous levels of heterogeneity within a few passages (11, 34). Here we saw that the modified Sia are retained in the Golgi of HEK-293 and A549 cells, although an occasional cell displayed these modified Sia on the surface. It is not clear why these modified Sia are localized in the Golgi, although perhaps the modifications could block the onward trafficking of glycoproteins to the cell surface. In contrast to the human cells lines, MDCK cells showed many cells expressing 7,9-*O*-Ac and 9-*O*-Ac on the cell surface, raising the possibility that trafficking could be cell-type or species specific. A summary of the synthesis and localization of 7,9-*O*- and 9-*O*-Ac is shown in **Figure 7**.

**Figure 7.**
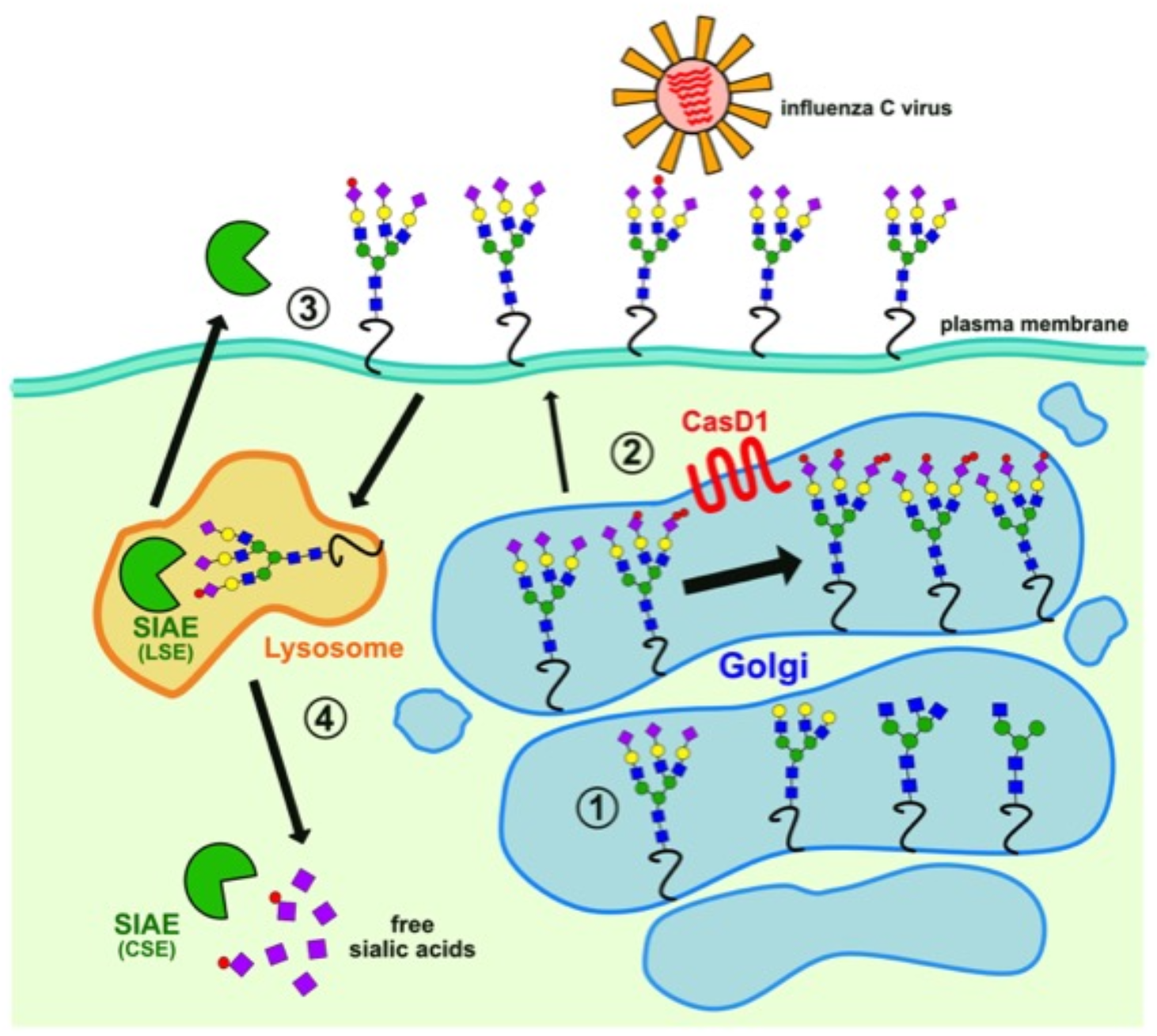
Summary of proposed 7,9-*O*- and 9-*O*-acetyl sialic acid production and trafficking in cells. (1) Sia (purple diamond) is added to the growing glycan chain in the Golgi by sialyltransferases using CMP-Neu5Ac or CMP-Neu5Gc substrates which are synthesized in the nucleus by the addition of cytodine monophosphate (CMP) to Neu5Ac or Neu5Gc, and are specifically imported into the Golgi. (2) Terminal Sia are modified by CasD1, adding one or two acetyl groups to form 9-*O*-Ac or 7,9-*O*-Ac, respectively (red circles). The majority of glycoproteins with these modifications are retained in the Golgi (large arrow) of many cells including the HEK-293 and A549 cells examined here, while only some, and mostly 9-*O*-Ac, are transported to the cell surface (small arrow). (3) Surface displayed *O*-acetyl Sia can interact with pathogens, cell receptors, or lectins. For example, influenza C virus uses 9-*O*-Ac as its receptor. Secreted forms of SIAE may also remove the *O*-acetyl modifications, altering these lectin-ligand interactions. (4) When glycoproteins are recycled from the cell surface, the lysosomal form of SIAE (LSE) can remove *O*-acetyl modifications from Sia. Free Sia are exported to the cytosol, where the cytosolic form of SIAE (CSE) can also remove any remaining *O*-acetyl modifications. Unmodified Sia can then be “activated” in the nucleus by the addition of CMP and transported to the Golgi for addition to new glycan chains.

To look more closely at the expression and roles of 7,9-*O*- and 9-*O*-Ac modifications and their effects on viral infections, we prepared glyco-engineered cell lines that lacked these modifications or that expressed higher levels of CasD1. A deletion and frame shift in CasD1 completely removed both 9-*O*- and 7,9-*O*-Ac expression, confirming this enzyme was responsible for creating both modifications, likely through addition to the C-7 position, from which it migrates to the C-9 position (5, 8, 9). Adding CasD1 back into the cells by plasmid transfection restored modified Sia expression, but none of the cell clones isolated showed universally higher levels of modified Sia synthesis and population heterogeneity was still seen. The CasD1-transfected HEK-293 cells showed the greatest increase, while for A549 and MDCK cells expression was similar to or lower than WT cells. However, even in the HEK-293 CasD1-OX cells, 9-*O*-Ac still only accounted for ~1.5% of total Sia and 7,9-O-Ac was not detected. This indicates that the levels of these modifications are not only controlled by the expression of CasD1, and could be regulated by other processes. Additionally, there is clearly a differential regulation of 7,9-*O*-Ac compared to 9-*O*-Ac, as over-expressing CasD1 did not lead to an increase in 7,9-*O*-Ac, as it was not detectable by HPLC and only very low fluorescence via staining with BCoV HE-Fc.

One candidate for control of these modifications is SIAE. When we knocked SIAE out of HEK-293 and A549 cells, we saw an increase in 9-*O*-Ac that was still retained in the Golgi but no increase in 7,9-*O*-Ac. Expressing CasD1 from a plasmid in ΔSIAE HEK-293 and A549 cells resulted in an additional small increase in 9-*O*-Ac that was still Golgi associated, but with little increase in expression on the cell surface. Similar to the CasD1-OX cells, no increase in 7,9-*O*-Ac was seen in either ΔSIAE and ΔSIAE+CasD1 cell lines by HPLC analysis. Both ΔSIAE+CasD1 lines had altered growth rates compared to WT cells: HEK-293 ΔSIAE+CasD1 had delayed growth rates, while A549 ΔSIAE+CasD1 had increased growth rates. This suggests that dysregulation of SIAE and CasD1 by gene manipulation affects cell metabolism and growth in a cell-type specific manner, possibly through the build-up of glycoproteins in the Golgi or through dysregulation of the sialic acid recycling pathway. These affects on cell growth could have implications for cancer and organismal development, as these variant Sia are involved in both these processes (15–17, 20–23, 41). Further research is needed to determine how 7,9-*O*- and 9-*O*-Ac expression is regulated, how they are transported within the ER and Golgi, and their roles in cell growth. Of particular interest will be disentangling the individual regulation of 7,9-*O*-Ac from 9-*O*-Ac, as their does seem to be differences in how they are regulated both in our cells and in previously reported expression in animal tissues (10, 11).

Studies with two strains of IAV showed no differences in infection efficiency in WT HEK-293 or MDCK-NBL2 cells compared to their ΔCasD1 or CasD1-OX variants. This is unsurprising, as >95% of the Sia is un-acetylated Neu5Ac which can be utilized by IAV as a receptor for binding and entry. However, the 1-2% of 9-*O*-Ac on the surface of WT MDCK-NBL2 cells was sufficient for ICV virus binding and entry, and that cell susceptibility was lost when CasD1 was inactivated. Similar results are likely seen for other viruses that use these modified Sia as a receptor, including human coronaviruses OC43 and HKU1 (42).

In summary, we have shown that these modifications are present in different cell lines sourced from different species of animals, but they make up a small minority of the total Sia present. In addition, these modifications vary considerably in their localization and have an inherent heterogeneity within cell populations. While the presence of both 7,9-*O*- and 9-*O*-Ac were dependent on the activity of CasD1, the relative proportions, levels of expression, and localization appear to be controlled by more complex mechanisms than simply the expression of CasD1 and SIAE. How this regulated expression affects cell homeostasis is unknown, but it is likely relevant during development, immune responses, and in cancers that show dysregulation of 7,9-*O*- and 9-*O*-Ac expression. For viruses such as ICV that rely on 9-*O*-Ac for infection the low levels seen on cell surfaces are sufficient for infection. While these modifications are present at levels too low to affect IAV, they are expressed at much higher levels in mucosal tissues and in secreted mucins of many animals and tissues, which may provide a more effective barrier (10, 11, 43–45). In future work we are examining these processes for mucus and other sources in different animals.

## MATERIALS AND METHODS

### Cells and virus

HEK-293, A549, MDCK-NBL2, MDCK type I, and MDCK type II cells were grown in Dulbecco’s Modified Eagle Medium (DMEM) with 5% fetal bovine serum and 50 μg/ml gentamicin. HEK-293, A549, and MDCK-NBL2 cells were obtained from ATCC. MDCK type I and type II cells were gifts from Dr. William Young (University of Kentucky) (31, 33, 46, 47). SiNEC and SiTEC cells were gifts from Dr. Stacey Shultz-Cherry (St. Jude’s Children Hospital). Influenza A virus strains pH1N1 (A/California/04/2009) and Victoria H3N2 (A/Victoria/361/2011) were rescued from reverse genetics plasmids using established protocols (48, 49). Rescued viruses were grown to low passage on MDCK-NBL2 cells using infection media containing DMEM, 0.03% BSA, and 1 μg/ml TPCK-treated trypsin. Influenza C virus C/Ann Arbor/1/50 and C/Taylor/1233/1947 were a gift from Dr. Andrew Pekosz (Johns Hopkins University) and were grown on MDCK-NBL2 cells using infection media containing DMEM, 0.03% BSA, and 5 μg/ml TPCK-treated trypsin.

### Immunofluorescence microscopy and flow cytometry

Cells were stained for using probes derived from viral hemagglutinin esterase proteins fused to human IgG1 Fc (HE-Fc). The porcine torovirus strain 4 HE-Fc (PToV HE-Fc) primarily recognizes 9-*O*-Ac and the bovine coronavirus Mebus strain HE-Fc (BCoV HE-Fc) recognizes 7,9-*O*-Ac and shows low levels of binding to 9-*O*-Ac (10, 11). For immunofluorescence microscopy, cells were seeded onto glass coverslips and left overnight at 37°C and 5%CO_2_. Coverslips were fixed in 4% paraformaldehyde (PFA) for 15 min. Coverslips were incubated with Carbo-Free Blocking Solution (Vector Laboratories) for 1 hr at room temperature, with optional permeabilization with 0.001% Tween-20. To stain, HE-Fc probes were pre-complexed with Alexa-488 labeled anti-human IgG antibody for 1 hr at 4°C then diluted in Carbo-Free blocking solution to a final concentration of 5 μg/ml HE-Fc and 1:500 of secondary antibody. Cells were stained with HE-Fc/anti-IgG complex for 1 hr at room temperature. Coverslips were mounted using Prolong Antifade-Gold with DAPI (Invitrogen). Cell were imaged using a Nikon TE300 fluorescent microscope. For flow cytometry, cells were seeded onto non-adherent cell culture dishes and left overnight at 37°C and 5% CO_2_. Cells were collected using ice-cold PBS (HEK-293 and A549 cells) or Accutase (Sigma, MDCK cells) to retain surface glycans, then fixed in 4%PFA for 15 min. Cells were blocked as above. HE-Fc probes were prepared as above with final concentrations of 5 μg/ml HE-Fc probe and 1:1200 of anti-IgG. A Millipore Guava EasyCyte Plus flow cyotometer (EMD Millipore, Billerica, MA) was used to collect data, analysis using FlowJo software (TreeStar, Ashland, OR). Statistical analyses were performed in PRISM software (GraphPad, version 8).

### Cell line mutations and characterization

Two methods for utilizing CRISPR-Cas9 were used. For A549 and MDCK cells, paired Cas9 plasmids (PX459, Addgene plasmid #62988) targeted adjacent sites in early exons of CasD1 as diagrammed in Figure 4A. Plasmids were transfected using TransIT-X2 (Mirus Bio LLC) (8). For HEK-293 cells, nickase Cas9 plasmids (PX462, Addgene plasmid #62987) were used instead. Transfected cells were selected with puromycin and single cell clones screened with PToV-P4 HE-Fc to identify non-staining variants. Edited sequences were confirmed by PCR amplification of the targeted regions, and sequencing the PCR product for each allele. Knock-out cell lines were used to prepare over-expression cell lines by transfection of a pcDNA3.1(-) plasmid expressing the complete human CasD1 cDNA open reading frame synthesized by Bio Basic (Markham, Ontario, Canada). Transfected cells were selected with G418 and single cell clones screened by staining with PToV-P4 HE-Fc to identify 9-*O*-Ac positive cell lines. Editing of the SIAE gene followed a similar protocol for CRISPR-Cas9 as above, and the gene regions targeted are shown in Figure 5A. After transfection and selection, cells were cloned and single-cell clones were screened by direct PCR amplification of the target gene region and analysis of the PCR product size for the edited form of both alleles. Full sequencing of each allele and qPCR were performed to confirm deletion of the gene.

### Quantification of Sia variants

The Sia composition of cells were determined by incubating with 2M acetic acid at 80°C for 3 hr, filtration through a Microcon 10 kD centrifugal filter (Millipore), and drying in a SpeedVac vacuum concentrator. Released Sia were derivatized with 1,2-diamino-4, 5-methylenedioxybenzene (DMB, Sigma Aldrich) for 2.5 hr at 50°C (35). HPLC analysis was performed using a Dionex UltiMate 3000 system with an Acclaim C18 column (ThermoFisher) under isocratic elution in 7%methanol, 7% acetonitrile, and 86% water. Sia standards included bovine sub-maxillary mucin and commercial standards for Neu5Ac and Neu5Gc (Sigma Aldrich). Statistical analyses were performed in PRISM software (GraphPad, version 8).

### Characterization of A549 conditioned media

Conditioned media from A549 cells was prepared by washing a fully confluent flask of cells to remove any serum, and allowing the cells to grow in serum-free media for 5-7 days. Conditioned media was collected, dialyzed with three volumes of PBS, and concentrated using a 30 kD centrifugal filter (Pall Corporation). Protein concentration was determined using a Qubit 4 fluorometer (Invitrogen). To determine Muc5B presence, two-fold dilutions of conditioned media were compared to purified human Muc5B using a 8% SDS-PAGE gel and probed with an anti-human Muc5B antibody (both purified protein and antibody were gifts from Dr. Stefan Ruhl, University of Buffalo). Conditioned media was analyzed for total Sia using the HPLC methods listed above.

### qPCR of SIAE and CasD1 expression

RNA from cells was extracted using EZNA Total RNA Kit I (Omega Bio-Tek) and cDNA was synthesized using SuperScript II Reverse Transcriptase (Invitrogen) standard protocol using oligo(dT)_12-18_ primers (Invitrogen). CasD1 and SIAE specific primers were designed using Geneious (Biomatters, Ltd.). For CasD1, adequate primers could not be targeted around the CRISPR-Cas9 edit site due to high G/C content. qPCR was performed on a Applied Biosystems StepOnePlus Real-Time PCR System using Fast SYBR Green (Bio-Rad). Data was analyzed using StepOne software (Applied Biosystems, version 2.1) and PRISM statistical analysis software (GraphPad, version 8).

### Virus infection assays

Influenza A viruses, cells were seeded on coverslips and inoculated with an MOI of 0.1. Coverslips of inoculated cells were fixed at 24 hrs and stained for virus using an anti-NP antibody and co-stained with DAPI. Percent-infected cells were determined by imaging coverslips and counting infected cells per field. Statistical analyses were performed in PRISM software (GraphPad, version 8). For influenza C, cells were inoculated for 48 hrs. Cells were then fixed and stained using a mouse anti-influenza C antibody (a gift from Dr. Peter Palese, Icahn School of Medicine at Mount Sinai). Cells were imaged using a Nikon TE300 fluorescent microscope.

## ACKNOWLEDGEMENTS

We thank Wendy Weichert for expert technical support. Brynn Lawrence for HA-Fc production.

---

**Supplemental Figure 1.**
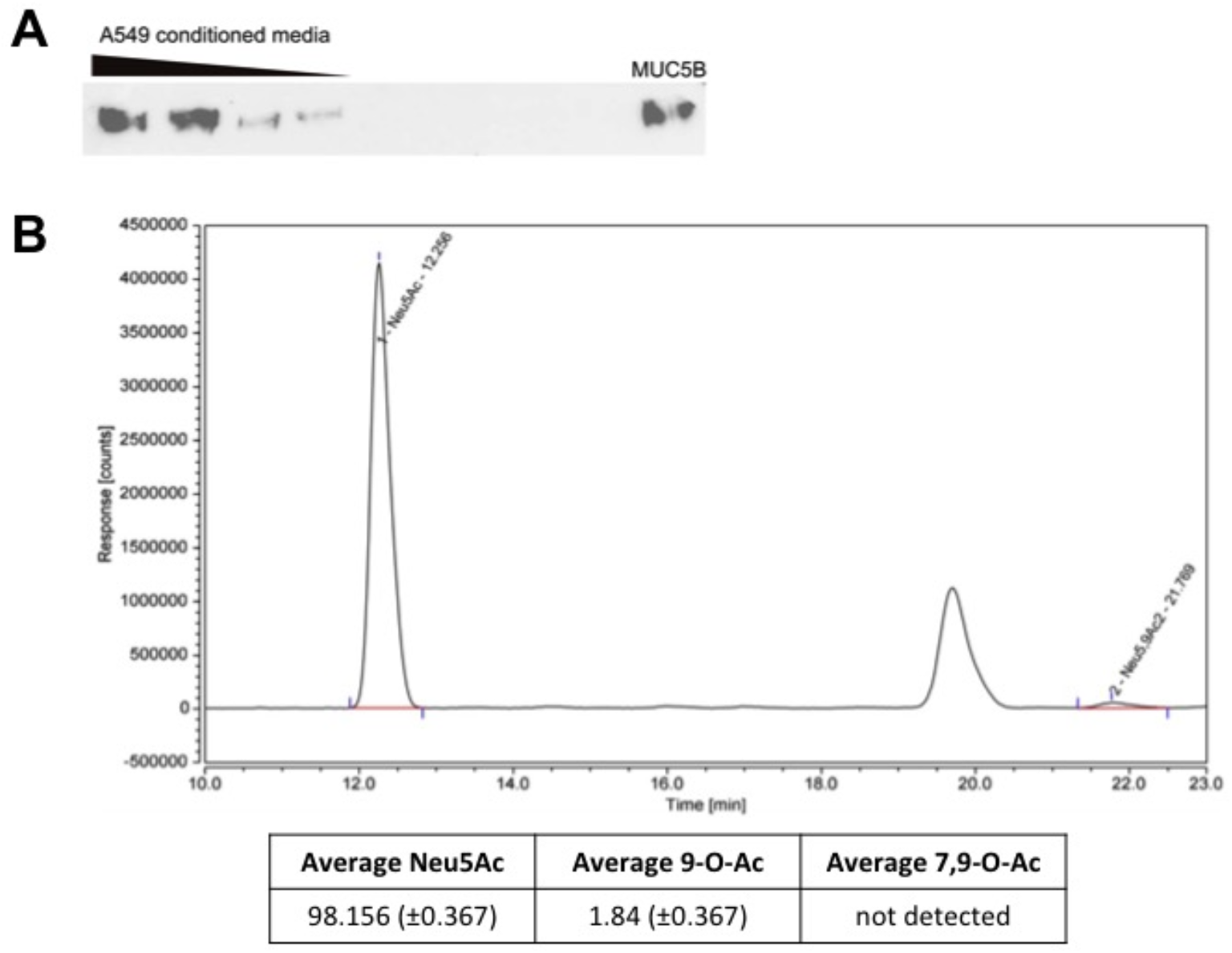
A549 conditioned media was analyzed for the presence of secreted mucins and total sialic acid content. A) Conditioned media from A549 was concentrated using a 30kd filter column and then titrated on a western blot for Muc5B expression compared to a purified human Muc5B (a gift from Stefan Ruhl, University of Buffalo). B) A representative chromatogram of total sialic acid collected from A549 conditioned media using HPLC analysis. The percent of different sialic acid forms found in total sialic acid collected is summarized in the table.

**Supplemental Figure 2.**
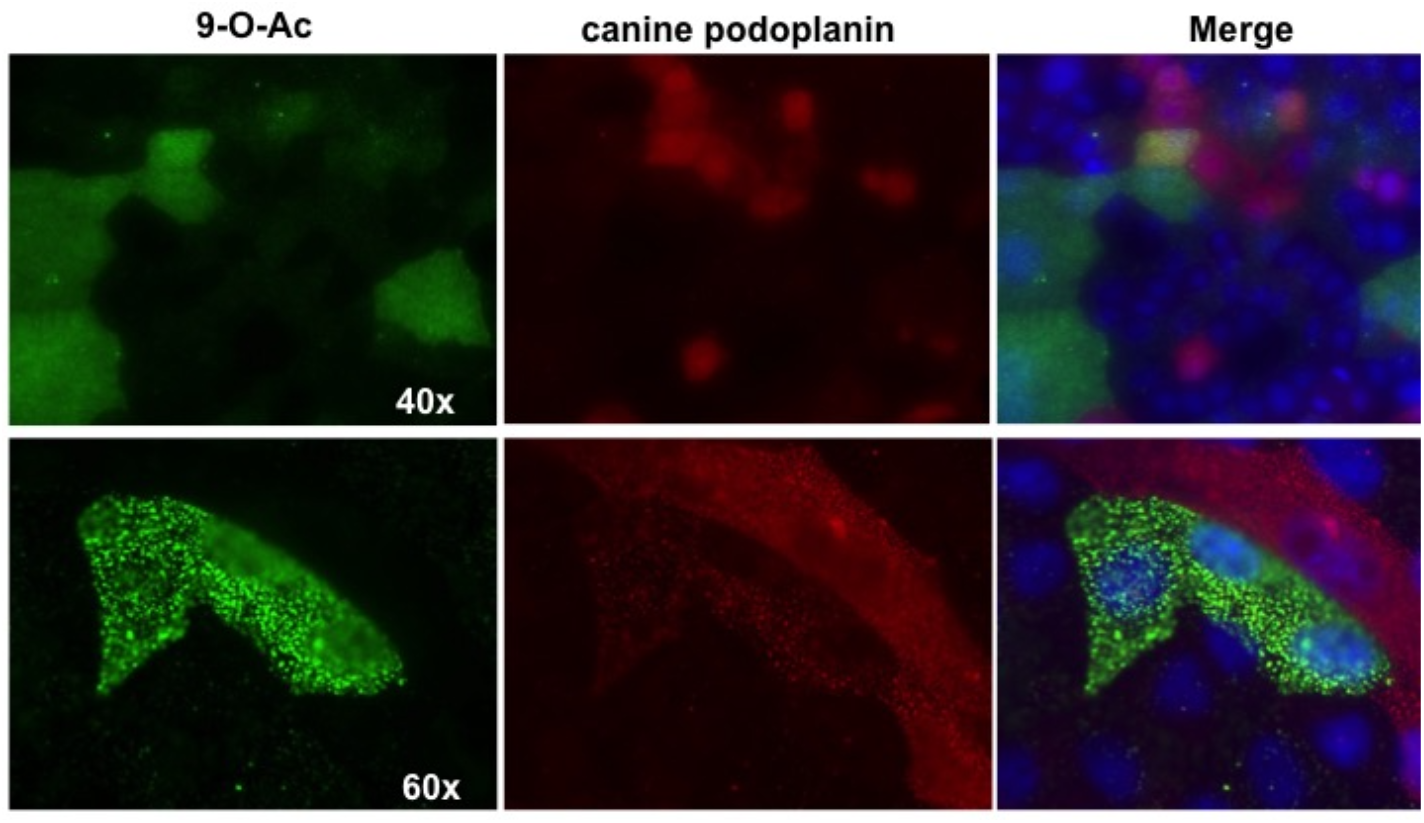
MDCK-NBL2 cells were co-stained for the presence of 9-*O*-Ac using PToV HE-Fc (green) and canine podoplanin (red) using an anti-podoplanin antibody (courtesy of Dr. Yukinari Kato, Tohoku University). Cells were imaged at 40× and 60× magnification as indicated.

**Supplemental Figure 3.**
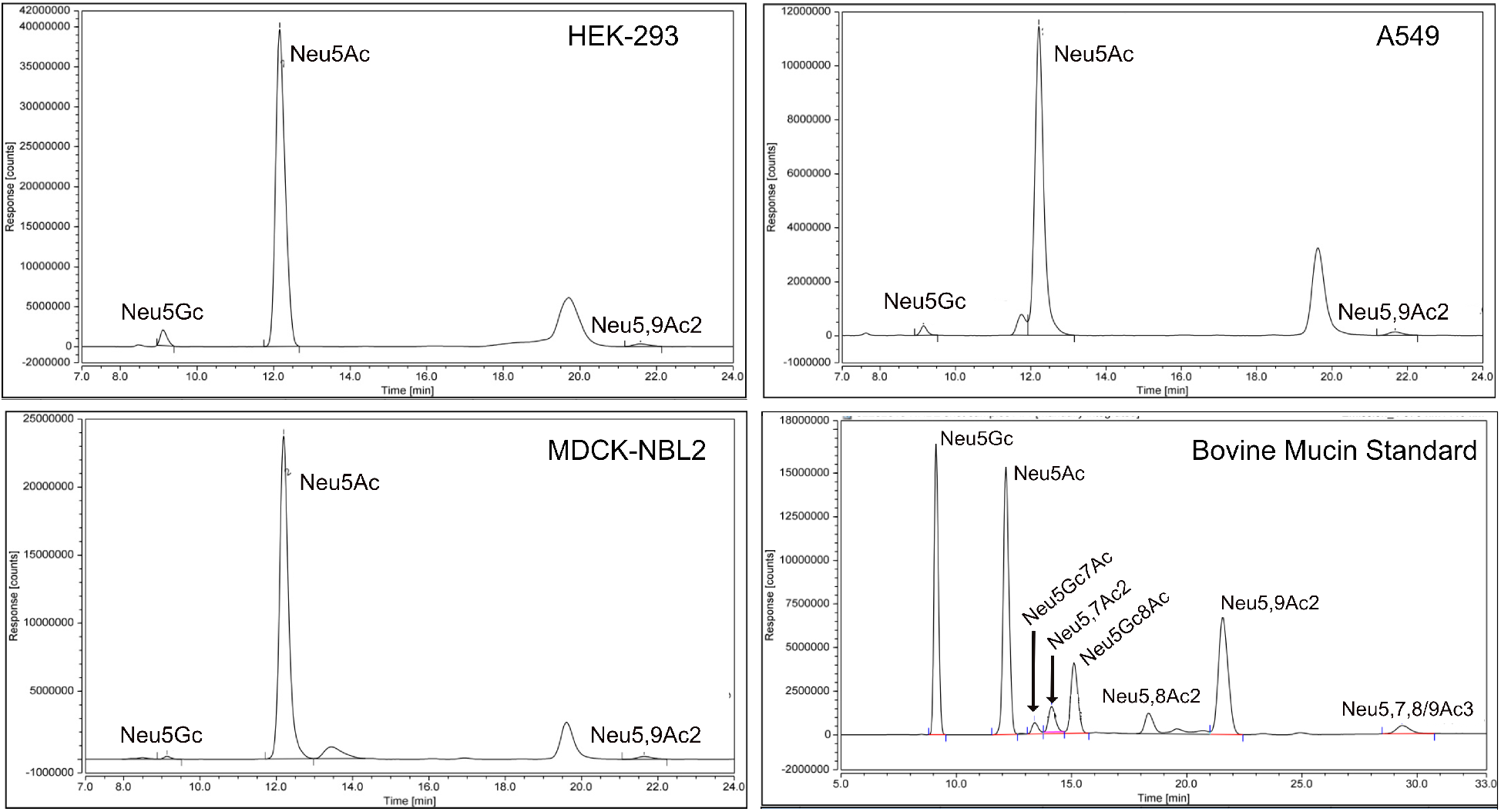
HPLC chromatograms showing wild-type HEK-293 cells, A549 cells, MDCK-NLB2 cells, and bovine mucin standard of *O*-acetyl modifications. Total sialic acids were collected from cell lines and standards via mild acid hydrolysis. Neu5Gc on cells is likely derived from the fetal bovine serum used in the growth media, which is taken up by cells and displayed on the cell surface. Humans and canines do not have a functional CMAH gene to synthesize Neu5Gc endogenously.

